# The subiculum encodes environmental geometry

**DOI:** 10.1101/2023.05.07.539721

**Authors:** Yanjun Sun, Douglas A Nitz, Xiangmin Xu, Lisa M Giocomo

## Abstract

Corners are a cardinal feature of many of the complex environmental geometries found in the natural world but the neural substrates that could underlie the perception of corners remain elusive. Here we show that the dorsal subiculum contains neurons that encode corners across environmental geometries in an allocentric reference frame. Corner cells changed their activity to reflect concave corner angles, wall height and the degree of wall intersection. A separate population of subicular neurons encoded convex corners. Both concave and convex corner cells were non-overlapping with subicular neurons that encoded environmental boundaries, suggesting that the subiculum contains the geometric information needed to re-construct the shape and layout of naturalistic spatial environments.

**One Sentence Summary:** Separate neural populations in the subiculum encode concave and convex environmental corners.

## Main Text

Animals in the natural world constantly encounter complex and non-symmetric landscapes, from networks of tree branches to winding burrow tunnels. Successful navigation requires understanding multiple features of these landscapes, including boundaries, discrete landmarks, and corners. Humans and other mammalian species use a combination of these features, which shape the overall geometric structure of the environment, to navigate (*1–9*), and neurons that contribute to building a ‘cognitive map’ for an environment, including hippocampal place cells (*10, 11*) and medial entorhinal grid cells (*12*), integrate information from these features (*13–21*). While the single-cell neural representations of environmental boundaries and discrete landmarks in both allocentric (world-centered) and egocentric (self-centered) reference frames has been well-documented in the hippocampus and surrounding parahippocampal cortices (*22–34*), whether and how the brain encodes environmental corners remains largely unexplored. One brain region that could play a role in encoding environmental corners is the subiculum, a major output structure of the hippocampus that receives highly convergent inputs from both the hippocampal sub-region CA1 and the medial entorhinal cortex (*35, 36*). Prior work has demonstrated that neurons in the subiculum encode the location of environmental boundaries and objects in an allocentric reference frame, as well as the axis of travel in multi-path environments (*22, 32, 37, 38*). Here, we describe single-cell neural representations for concave and convex environmental corners in the dorsal subiculum, which reside interspersed with single-cell neural representations for environmental boundaries. These results raise the possibility that the subiculum is ideally suited to re-construct the geometric shape and layout of an environment.

### Subiculum neurons encode corners across environmental geometries

To record from large numbers of neurons in the subiculum, we performed in vivo calcium imaging using a single photon (1P) miniscope in freely behaving mice (Fig 1A, B). We primarily used Camk2a-Cre; Ai163 (*39*) transgenic mice, which exhibited stable GCaMP6s expression in subiculum pyramidal neurons (Fig S1A) and thus facilitated longitudinal tracking of individual neurons across experiments (Fig. 1C) (*40*). Calcium signals were extracted using a CNMF based method and binarized into deconvolved spikes (Fig. 1D and Fig. S1B). We treated these deconvolved spikes as equivalent to electrophysiological spikes and for calculating spike rates in downstream analyses.

**Fig. 1.**
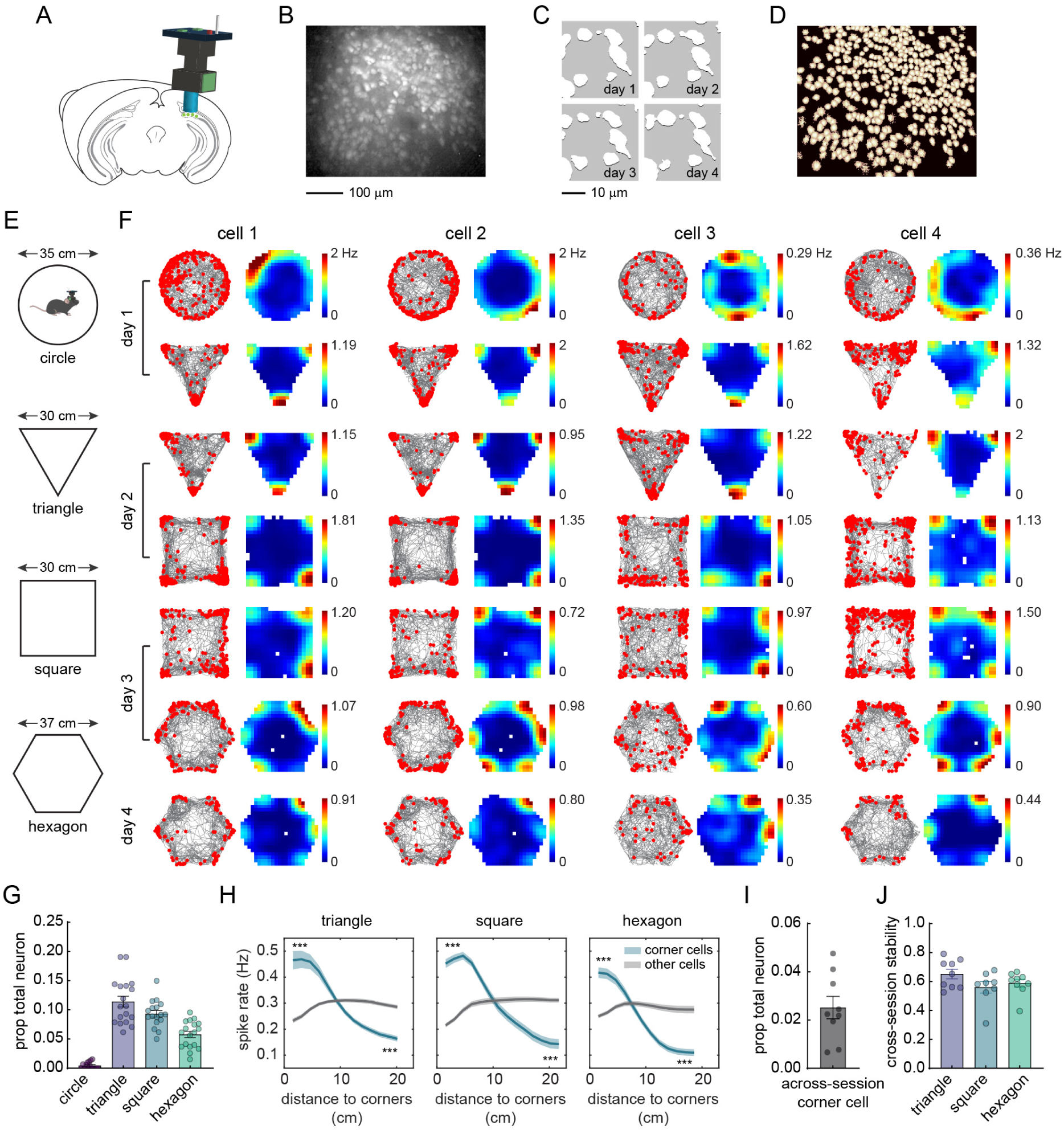
The subiculum contains neurons that exhibit corner-associated activity. **(A)** Schematic of calcium imaging in the subiculum using a miniscope. **(B)** Maximum intensity projections of subiculum imaging from a representative mouse. **(C)** An enlarged region of interest from (B), showing the persistent activity of the same neurons across days. An image filter was applied to better indicate the neurons (white). **(D)** Spatial footprints of the CNMF-E extracted neurons from (B). **(E)** Schematic of the open arena environment shapes mice explored over days. **(F)** Examples of corner-associated activity in four neurons from three different mice. Each column is a cell in which its activity was tracked across sessions and days. Raster plot (left) indicates extracted spikes (red dots) on top of the animal’s running trajectory (grey lines) and the spatial rate map (right) is color coded for maximum (red) and minimum (blue) values. **(G)** Proportion of neurons classified as corner cells in each environment shape. Each dot represents a session, with a maximum of two sessions per condition per mouse (n = 15, 18, 17, and 18 sessions for circle, triangle, square, and hexagon, respectively, from 9 mice). Histogram and error bars indicate mean ± standard error of the mean (SEM). **(H)** Positional spike rates plotted relative to the distance to the nearest environmental corner for the triangle, square and hexagon arenas. Statistical tests were conducted on the comparison between identified corner cells and other recorded neurons within ∼5 cm of the corners (head of the curve, averaged from the first 3 bins, 1 bin = 1.6 cm) and ∼5 cm of the environmental center (tail of the curve, averaged from the last 3 bins) (two-tailed paired t-test: all p < 0.0001, n = 18, 17, and 18 sessions for triangle, square, and hexagon, respectively, from 9 mice). Solid line: mean; Shaded area: SEM. **(I)** Proportion of cells classified as corner cells across sessions. Each dot represents a mouse (n = 9). Histogram and error bars indicate mean ± SEM. Neurons included here had to be classified as corner cells in at least one of the sessions in each non-circle condition. **(J)** Cross-session stability (Pearson’s correlation) of across-session corner cells for each environmental shape.

We placed the animals in one of four open field arenas, including a circle, an equilateral triangle, a square and a hexagon. One each day, we recorded subiculum neurons from two of these four different arenas (20 minutes per session) (Fig. 1E, F). Many subicular neurons exhibited place cell-like firing patterns that did not significantly change across the different geometric environments (Fig. S1C), as previously reported (*41, 42*). Moreover, subicular neurons provided significant information regarding the animal’s position in space, as indicated by a decoder (Fig. S2A, B). However, a subset of subicular neurons were active near the boundaries of the circle (Fig. 1F). Surprisingly, following the activity of these neurons in all the other non-circle environments revealed that they exhibited increased spike rates specifically at the corners of the environments (Fig. 1F). To ensure these neurons were anatomically located in the subiculum, we used a viral strategy to restrict GCaMP expression to the subiculum (Fig. S1D) and observed the same corner-associated neural activity (Fig. S1E).

To classify neurons that exhibited corner-specific activity patterns, we devised a corner score that measures how close a given spatial field is to the nearest corner (Fig. S1F). The score ranged from -1 for fields situated at the centroid of the arena, to +1 for fields perfectly located at a corner (Fig. S1F, Methods). We defined a corner cell as a cell with, i) a corner score greater than the 95^th^ percentile of a distribution of scores generated by shuffling the spike times along the animal’s trajectory (1000 shuffles) shuffled score (Fig. S1G-I) and, ii) a distance between any two fields greater than half the distance between the corner and centroid of the environment (Fig. S1J). Using this definition, we classified 11.4 ± 0.9% (mean ± SEM, n = 18 sessions from 9 mice, maximum of 2 sessions per mouse) of neurons as corner cells in the triangle, 9.3 ± 0.6% in the square (n = 17 sessions), and 5.8 ± 0.5% in the hexagon (n = 18 sessions) (Fig. 1G). Plotting the data for each mouse revealed very similar percentages (Fig. S1K). Notably, this method classified almost no neurons as corner cells in the circle (0.45 ± 0.14%, n = 15) when four equally spaced, arbitrary points were assigned as the ‘corners’ for the environment (Fig. 1G).

Verifying that neurons classified as corner cells encode spatial locations near corners, a decoder with classified corner cells removed resulted in higher decoding errors near the corners than at the center of the environment, compared to a full decoder (Fig. S2C, D). By plotting the spike rate for each bin on the rate map as a function of the distance to the nearest corner, we found that corner cells showed a significantly higher spike rate around (tested within 5 cm) the corners and a significantly lower spike rate around the centroid, compared to non-corner cells from the same animal (Fig. 1H). Importantly, as measured by the corrected peak spike rate at each corner, which accounted for the animals’ occupancy and movement (Fig. S2E, methods), corner cell population activity was not biased towards encoding specific corners (Fig. S2F). Finally, across all non-circle environmental geometries, 2.5 ± 0.5% of neurons were consistently classified as corner cells (Fig. 1F, I). These neurons, referred to as ‘across-session corner cells’, exhibited stable corner-associated activity in all environments (Fig. 1J) (mean cross-session stability from 0.56 – 0.65, Pearson’s correlation). Of note, the neural population classified as corner cells in one environment still exhibited activity at corners in later sessions/conditions in which they were not classified as corner cells (Fig. S2G, H), indicating corner activity generally persisted across different geometries when considering the subiculum neural population rather than only single cells classified based on their corner score.

### The activity of corner cells is specific to environmental corners

To investigate the degree to which corner cells specifically encode environmental corners, we considered three properties that comprise a corner: 1) the angle of the corner, 2) the height of the walls, and 3) the connection between two walls. First, we imaged as animals explored two asymmetric environments: a right triangle (30-60-90° corners) or a trapezoid (55-90-125° corners) (Fig. 2A). In these asymmetric environments, corner cells composed 5.4 ± 0.3% and 3.3 ± 0.3% of all neurons recorded in the right triangle and the trapezoid, respectively (Fig. 2B, C; n = 16 sessions from 8 mice). In the right triangle, corner cell peak spike rates were significantly higher for the 30° (2.08 ± 0.13, mean ± SEM) corner compared to the 60° (1.57 ± 0.09) and 90° (1.70 ± 0.11) corners, but did not differ between the 60° and 90° corners (Fig. 2B, D). To rule out the possibility that this was due to the limited angular range of these acute angles, we further compared the peak spike rates at the corners of the trapezoid (n = 14 sessions from 8 mice). Interestingly, the peak spike rates of corner cells increased from 125° (1.35 ± 0.07, mean ± SEM) to 90° (1.76 ± 0.08) to 55° (2.00 ± 0.07) (Fig. 2E). We also compared the peak spike rates at the corners using the aforementioned across-session corner cells in the triangle (60°, 1.62 ± 0.15, mean ± SEM), square (90°, 1.52 ± 0.07) and hexagon (120°, 1.41 ± 0.13). However, spike rates did not vary relative to the corner angle across different environments (Fig. 2F), indicating the spike rates of corner cells only differed when a single environment contained different corner angles. Together, these results suggest that corner cells encode information regarding corner angles within an asymmetric environment.

**Fig. 2.**
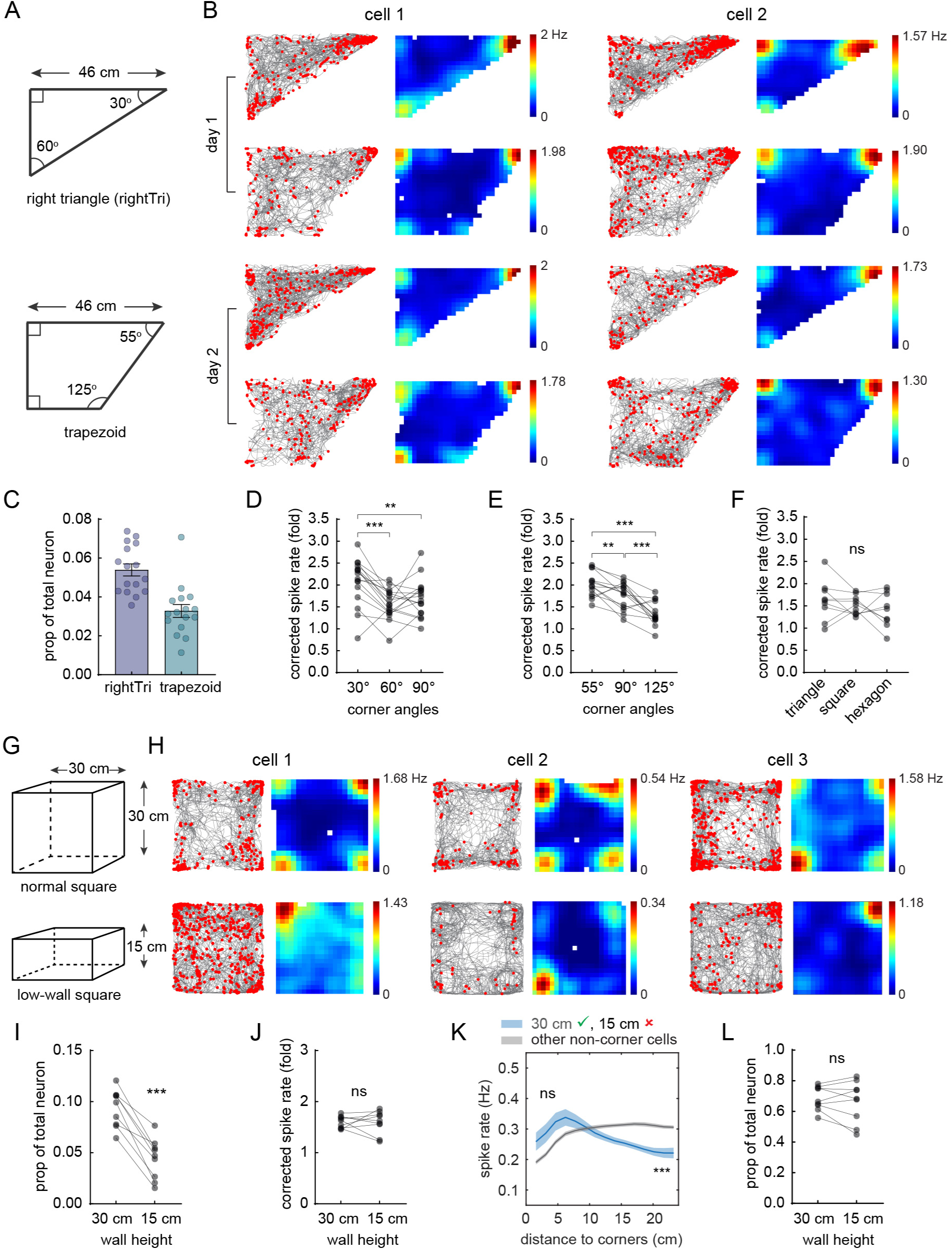
Corner cell coding is sensitive to corner angle and wall height. **(A)** Schematic of the open arena environment shapes (a 30-60-90 degree right triangle and a trapezoid). **(B)** Raster plots and the corresponding rate maps of two corner cells from two different mice (two sessions for each condition), plotted as in Fig. 1F. Each column is a cell in which its activity was tracked across sessions and days. **(C)** Proportion of neurons classified as corner cells in each environment shape. Each dot represents a session (two sessions per condition per mouse, n = 16 sessions from 8 mice). Histogram and error bars indicate mean ± SEM. **(D)** Corrected peak spike rates of corner cells within 10 cm of the corners of the right triangle (see Methods and Fig. S2), (repeated measures ANOVA: p = 0.0006; two-tailed paired t-test: 30° vs. 60°: t(15) = 4.55, p = 0.0004; 30° vs. 90°: t(15) = 2.98, p = 0.0093; 60° vs. 90°: t(15) = 1.12, p = 0.28; n = 16 sessions). **(E)** Corrected peak spike rates of corner cells within 10 cm of the corners of the trapezoid (repeated measures ANOVA: p < 0.0001; two-tailed paired t-test: 55° vs. 90°: t(13) = 3.05, p = 0.0094; 55° vs. 125°: t(13) = 7.51, p < 0.0001; 90° vs. 125°: t(13) = 4.20, p = 0.0010; n = 14 sessions. Note, two of the 16 sessions from two mice did not meet the data inclusion criteria, i.e., number of corner cells ≥ 5, see Methods). **(F)** Corrected peak spike rates of across-session corner cells at the corners of the triangle, square and hexagon (repeated measures ANOVA: p = 0.26; n = 9 mice). **(G)** Schematic of the open arena squares with different wall heights. The 30 cm wall height matched the wall height of environments used in Fig 1E and S3E. **(H)** Raster plots and rate maps of three corner cells from three different mice, plotted as in (B). **(I)** Proportion of neurons classified as corner cells in the 30 vs. 15 cm wall height square (two-tailed paired t-test: t(8) = 8.18, p < 0.0001; n = 9 mice. Data from each mouse was averaged across 1-2 sessions). **(J)** Corrected peak spike rates of corner cells within 10 cm of the corners defined in the 30 vs. 15 cm wall condition (two-tailed paired t-test: t(8) = 0.090, p = 0.93). **(K)** Positional spike rates plotted relative to the distance to the nearest corner in the 15 cm wall square condition. Blue curve indicates neurons that were a corner cell in the 30 cm wall arena (green check mark) but not in the 15 cm wall arena (red cross). Grey curve indicates other non-corner cells in the 15 cm wall arena. Statistical tests were performed as in Fig. 1H (two-tailed paired t-test: head of the curve: t(8) = 2.20, p = 0.06; tail of the curve: t(8) = -5.99, p = 0.0003; n = 9 mice) **(L)** Same as I, but for comparisons of place cells (i.e., all neurons that spatially tuned) in the 30 vs. 15 cm wall arenas (two-tailed paired t-test: t(8) = 0.84, p = 0.43; n = 9 mice). For all panels, ** p < 0.01; *** p < 0.001; ns: not significant.

Next, we imaged as animals explored the normal square environment (as in Fig. 1) with 30 cm high walls, followed by a low-wall square environment with 15 cm high walls (Fig. 2G, H). Quantitative analysis revealed the proportion of corner cells significantly decreased from the normal (9.4 ± 0.6%, mean ± SEM) to the low-wall square (4.4 ± 0.7%) (Fig. 2I). The remaining corner cells in the low-wall square had corner spike rates similar to their corner spike rates in the normal square (1.60 ± 0.04 vs. 1.60 ± 0.08; n = 9 mice) (Fig. 2J). However, the spike rates near the corners for neurons classified as corner cells in the normal square but not in the low-wall square did not differ from the spike rates of non-corner cells in the low-wall square (Fig. 2K). This was a more degraded corner cell representation when compared to a similar analysis on corner cells across two normal square sessions (Fig. S2H). Finally, in comparison to corner cells, the proportion of subiculum place cells did not change between the normal (68.5 ± 2.6%) and low-wall squares (66.4 ± 4.5%) (Fig. 2L). Together these results indicate the tuning of corner cells is sensitive to the height of the walls that constitute the corner.

Finally, we imaged as animals explored a large square environment in which we inserted a discrete corner and gradually separated its two connected walls (1.5, 3, or 6 cm separation) (Fig. 3A). We identified corner cells in the baseline session and tracked their activity across all manipulations (Fig. 3B). Despite the insertion of the discrete corner, corner cells identified in baseline did not change their average peak spike rates across other conditions at the corners of the square environment (Fig. 3B, C). Upon the insertion of the discrete corner, corner cells developed a new field near the inserted corner (Fig. 3B). Interestingly, as the distance between the walls of the discrete corner increased, the peak spike rate of corner cells at that corner gradually decreased (Fig. 3D). Even at the largest gap of 6 cm, however, corner cell peak spike rate at the discrete corner was still significantly higher than at baseline (1.11 ± 0.13 vs. 0.42 ± 0.05, mean ± SEM) (Fig. 3D), indicating that the animal may still perceive the inserted walls as a corner. Furthermore, we observed no difference in spike rate between the 0 cm (1.76 ± 0.15) and 1.5 cm (1.55 ± 0.19) gap, suggesting a physical connection between the walls was not necessary to evoke corner cell firing (Fig. 3D). Even so, the peak spike rates of corner cells at 3 cm (1.20 ± 0.08) and 6 cm (1.11 ± 0.13) gap were significantly attenuated compared to the 0 cm (1.76 ± 0.15) gap condition (Fig. 3D), suggesting corner cells are sensitive to the proximity of the walls that constitute the corner. In comparison, there was no effect at the inserted corner when we performed the same analyses using non-corner cells (Fig. 3E).

**Fig. 3.**
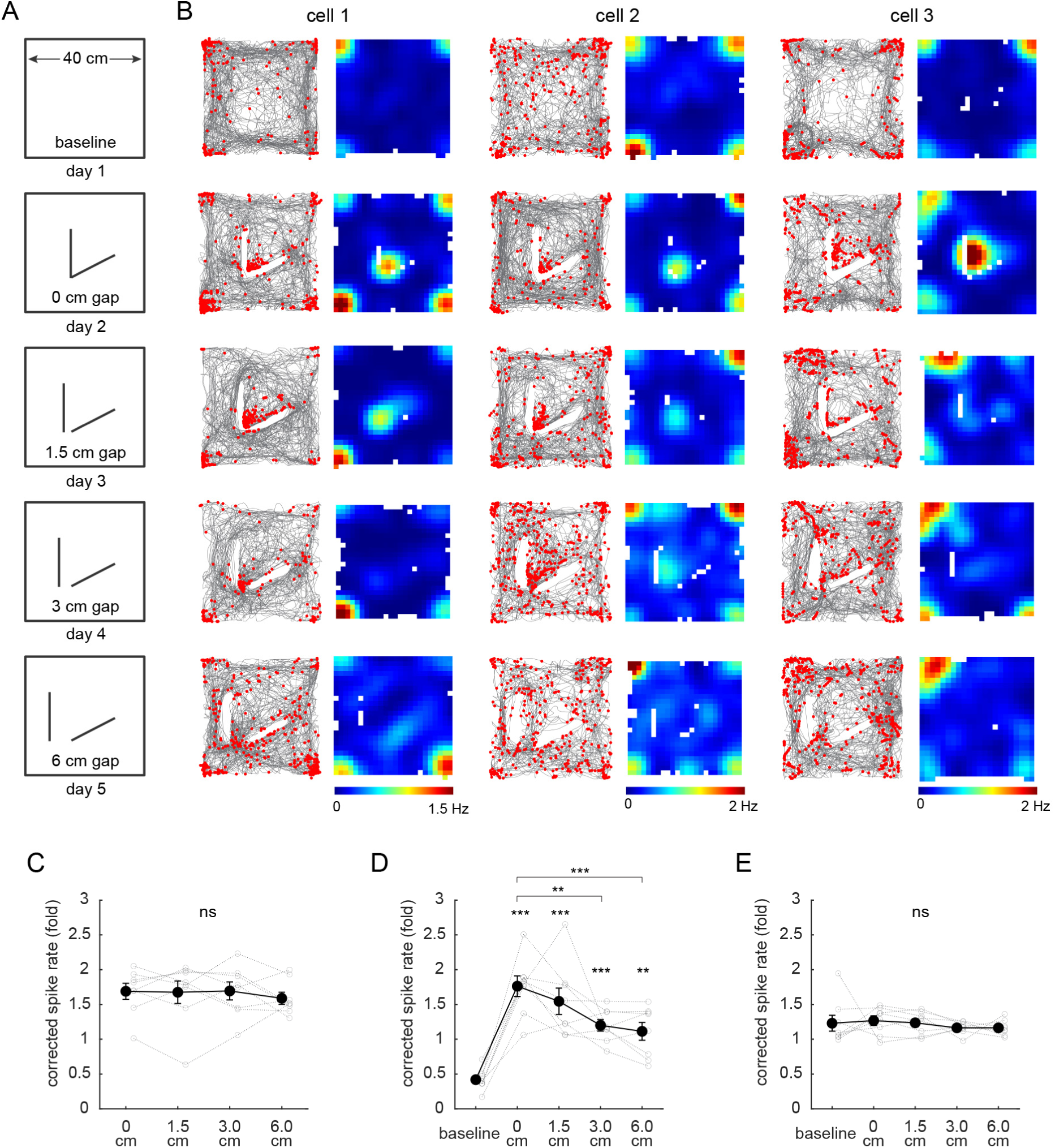
Corner cell coding is sensitive to the proximity of the walls that constitute a corner. **(A)** Schematic of the open field arena and sessions in which a discrete corner was inserted into the center of the environment. **(B)** Raster plots and the corresponding rate maps of three corner cells from three different mice, plotted as in Fig. 1F. Each column is a cell in which its activity was tracked across sessions. Note that rate maps for each cell were plotted to have the same color-coding scale for maximum (red) and minimum (blue) values. **(C)** Corrected peak spike rates of baseline-identified corner cells at the environmental corners (not the inserted corner) across non-baseline sessions (repeated measures ANOVA: p = 0.75; n = 8 sessions from 8 mice). Black dots represent mean ± SEM, while the grey lines represent each animal. **(D)** Corrected peak spike rates of baseline-identified corner cells at the inserted corner across all the sessions, plotted as in (C) (repeated measures ANOVA: p < 0.0001; two-tailed paired t-test: baseline vs. 0 cm: t(7) = 9.63, p < 0.0001; baseline vs. 1.5 cm: t(7) = 7.92, p < 0.0001; baseline vs. 3 cm: t(7) = 7.73, p < 0.0001; baseline vs. 6 cm: t(7) = 5.02, p = 0.0015; 0 cm vs. 1.5 cm: t(7) = 1.14, p = 0.29; 0 cm vs. 3 cm: t(7) = 4.77, p = 0.0020; 0 cm vs. 6 cm: t(7) = 6.01 p = 0.0005; n = 8 sessions from 8 mice). **(E)** Same as (D), but for non-corner cells (repeated measures ANOVA: p = 0.63; n = 8). For all panels, ** p < 0.01; *** p < 0.001; ns: not significant.

### Corner coding is not sensitive to non-geometric features and is driven by multimodal sensory inputs

We next investigated whether corner cells in the subiculum are sensitive to non-geometric features of a corner. To test this, we placed the animals in a shuttle box with two connected square compartments that differed in color and texture (Fig. S3A). Example corner cells showed increased spike rates uniformly across all the corners, regardless of the context, suggesting that color and texture did not significantly affect the activity of these cells (Fig. S3B). Quantitative analysis supported this observation, showing similar proportions of corner cells across the two contexts with comparable average peak spike rates at corners (Fig. S3C, D). These results suggest that corner cells in the subiculum primarily encode corner-associated geometric features, rather than non-geometric properties, such as colors and textures.

We then placed animals in a square arena in complete darkness. In this condition, the representations of corners by corner cells persisted, and the proportion of corner cells remained unchanged (Fig. S3E-G). Similarly, trimming the animals’ whiskers did not significantly affect the proportion of corner cells (Fig. S3E-G). There was also no difference in the peak spike rates of corner cells at the corners across all three conditions (Fig. S3H). In contrast, consistent with previous reports in the hippocampus (*43–45*), recording in darkness or with trimmed whiskers significantly decreased the number of place cells in the subiculum (Fig. S3I). Together, our results suggest that either visual or somatosensory inputs may be sufficient to drive the representation of corner cells in a familiar environment. One possibility is that these inputs are complementary, with each providing input that can be used to identify a corner, even when information from one sensory system is lacking.

### Subiculum neurons encode convex corners

If corner sensitivity in the subiculum plays a significant role in encoding environmental geometry, it would be reasonable to anticipate distinct coding for concave versus convex corners, as these qualitative distinctions are critical for defining geometry. We next examined whether corner coding in the subiculum extended to other corner geometries. We designed more complex environments that included both concave and convex corners. We imaged as animals explored a square and rectangle environment (concave corners, 30 minutes), followed by three environments with convex corners (convex-1, convex-2, convex-3) (Fig. 4A). First, we identified corner cells in the square and followed their activity across other environments. As in our prior experiments, we observed corner cells that increased their spike rate at the concave corners, but less so to the convex corners (Fig. 4B). Further investigation of neurons imaged in the convex-1 environment however, revealed a small sub-set of neurons that increased their spike rate specifically at the convex corners (Fig. 4C). By tracking the activity of these convex corner cells to the convex-2 and -3 environments, we further found that they responded to convex corners regardless of the location of the corners or the overall geometry of the environment (Fig. 4C). Compared to other cells in the subiculum from the same animal, convex corner cells showed a significantly higher spike rate around the convex corners as well as a significantly lower spike rate around the centroid (Fig. 4D) (n = 10 sessions from 10 mice). Tracking the activity of convex corner cells retrogradely to the square and rectangle environments, we observed that they had an overall lower spike rate compared to other subicular neurons (Fig. 4D). This low level of activity of convex corner cells in the absence of convex corners suggests they are convex corner specific. In environments with convex corners, the proportion of convex corner cells was 1.6 ± 0.2%, a slightly smaller proportion than concave corner cells identified in the square (3.6 ± 0.8%) and rectangle (4.1 ± 0.7%) in the same experiment (Fig. 4E). Concave and convex corner coding rarely occurred in the same neuron, suggesting convex corner cells are a separate neuronal population from concave corner cells (Fig. 4F). Anatomical analysis further suggested concave and convex corner cells were distributed in the subiculum in a salt and pepper pattern without clear clustering (Fig. 4G, H). Similar to concave corner cells, the activity of convex corner cells was not affected by non-geometric changes to the corners, showing consistent spike rates for the same corner before and after it was decorated with different colors and textures (Fig. S4A-C). However, unlike concave corner cells, convex corner cells showed comparable spike rates for corners at various angles in an asymmetric environment (315°: 2.01 ± 0.25, 270°: 2.12 ± 0.27 & 2.09 ± 0.11, and 225°: 2.18 ± 0.17, mean ± SEM) (Fig. 4B, I; Fig. S4D-F). These results demonstrate that concave and convex corner coding exists in the subiculum, likely as two distinct neuronal populations.

**Fig. 4.**
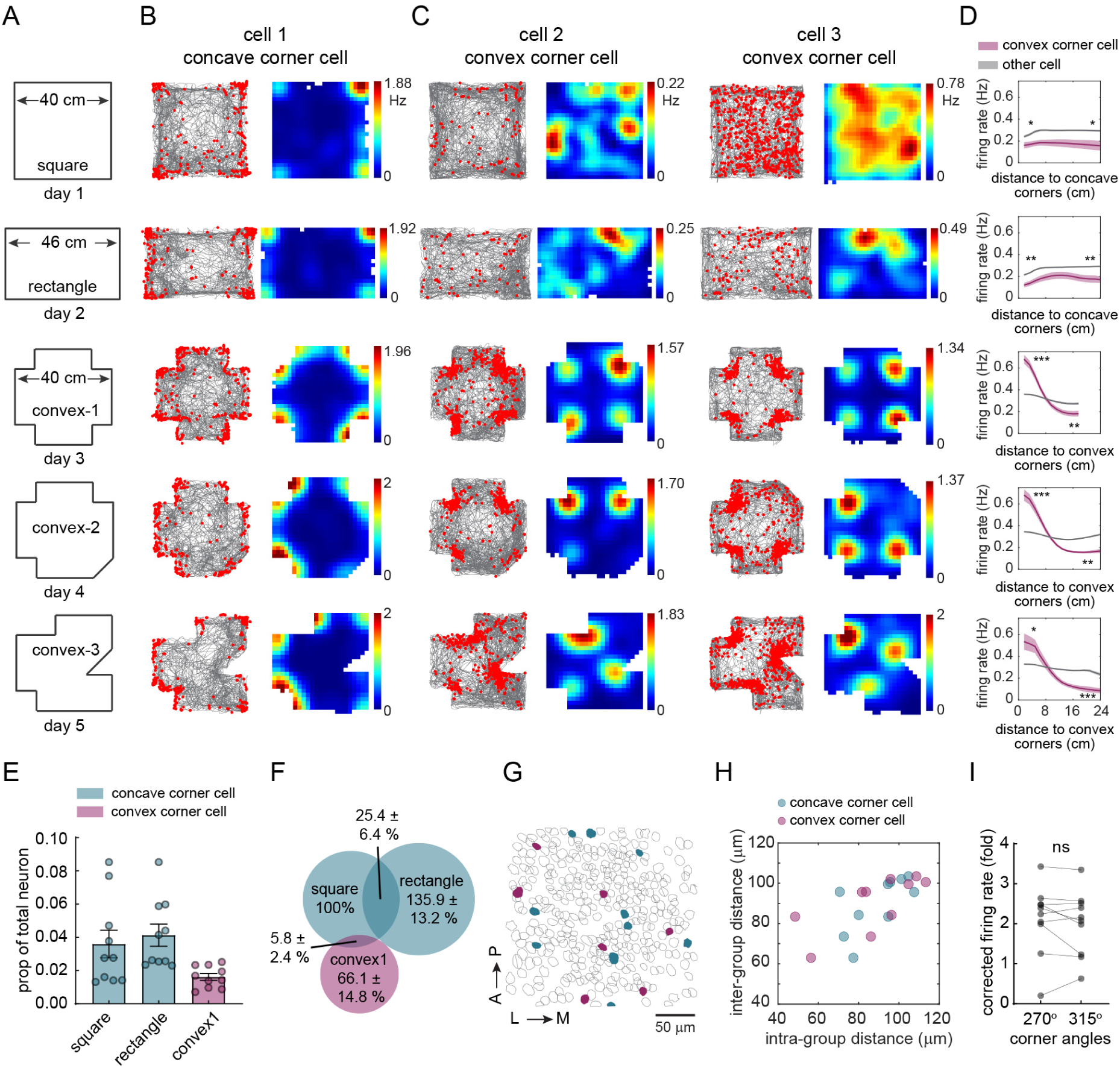
The subiculum contains neurons that exhibit convex corner-associated activity. **(A)** Schematic of the open field arenas that contained both concave and convex corners. **(B)** Raster plots and the corresponding rate maps of a representative concave corner cell with activity tracked across all the sessions, plotted as in Fig. 1F. **(C)** Raster plots and rate maps of two convex corner cells from two different mice. Each column is a cell in which its activity was tracked across sessions. **(D)** Positional spike rates plotted relative to the distance to the nearest convex corner for convex corner cells. Plots are organized as in (A). Data in the square and rectangle (top two plots) show positional spike rates of convex corner cells identified in the convext-1 environment and relative to the distance to the nearest concave corner. Statistical tests compared spike rates between convex corner cells and other recorded neurons within 5 cm of the corners (head of the curve) and 5 cm of the environment center (tail of the curve), as in Fig. 1H (two-tailed paired t-test: * p < 0.02, ** p ≤ 0.003, *** p ≤ 0.001; n = 10 sessions from 10 mice). **(E)** Proportion of neurons classified as concave corner cells in the square and rectangle, and convex corner cells from the convex-1 arena. Each dot represents a session (n = 10). Histogram and error bars indicate mean ± SEM. **(F)** Venn diagram showing the overlap between classified concave and convex corner cells. All numbers were normalized to the number of concave corner cells in square. **(G)** Anatomical locations of concave and convex corner cells from a representative mouse. Teal color represents concave corner cells from square, while purple color represents convex corner cells from the convex-1 arena. Unfilled grey circles represent non-corner cells. A: anterior; P: posterior; L: lateral; M: medial. **(H)** Pairwise intra- vs. inter-group anatomical distances for concave and convex corner cells (repeated measures ANOVA: p = 0.77; n = 10). **(I)** Corrected peak spike rates of convex corner cells (identified in the convex-3 arena) at the convex corners (270° vs. 315°) in the convex-3 arena (two-tailed paired t-test: t(9) = 0.99, p = 0.35; n = 10).

### Corner coding in the subiculum is allocentric

To determine whether the observed concave and convex corner coding is allocentric or egocentric, we trained a linear-non-linear Poisson (LN) model with behavioral variables including the animal’s allocentric position (P), head direction (H), running speed (S) and egocentric bearing to the nearest corner (E) (Fig. S5A-D). We used concave corner cells from the square environment (40 cm) and convex corner cells from the convex-1 environment for this analysis. For both concave and convex corner cells, the majority (i.e., 45 out of 71 LN classified concave corner cells and 14 out of 35 LN classified convex corner cells. Note, 34 concave and 14 convex corner cells could not be classified in the LN model) fell into the allocentric position only category (P), which means that adding variables did not improve the model performance (Fig. S5E-H). Only a small number of corner cells encoded head direction, running speed, or/and egocentric corner bearing (Fig. S5G-H), indicating corner coding in the subiculum is largely independent of modulation by the animal’s head direction, running speed and egocentric corner bearing. Furthermore, a decoder trained using only corner cells was able to predict the animal’s quadrant location significantly better than chance in the square environment (Fig. S5I-K). Our results suggest that concave and convex corner coding is primarily allocentric, a reference frame consistent with BVC and place cell coding in the subiculum (*22, 32, 37*).

### Corner coding differs from boundary coding in the subiculum

Finally, we examined the relationship between corner cells and previously reported boundary vector cells (BVCs) in the subiculum (*22*). We observed BVCs in the square (11.9 ± 0.9%, n = 10 session from 10 mice) and rectangle (7.6 ± 0.8%) environments (Fig. 5A, B). Tracking the activity of BVCs identified in the square environment revealed stable boundary coding across both concave and convex environments (Fig. 5A). We observed a small degree of overlap between BVCs and concave corner cells (Fig. 5C, D), reflective of neurons that were active at both corners and boundaries (Fig. 5D). However, we did not observe any overlap between BVCs and convex corner cells (Fig. 5C). Anatomically, BVCs and corner cells did not form distinct clusters but instead showed a salt-and-pepper distribution in the subiculum (Fig. 5E, F). Together, corner cells are a separate neuronal population from boundary vector cells in the subiculum.

**Fig. 5.**
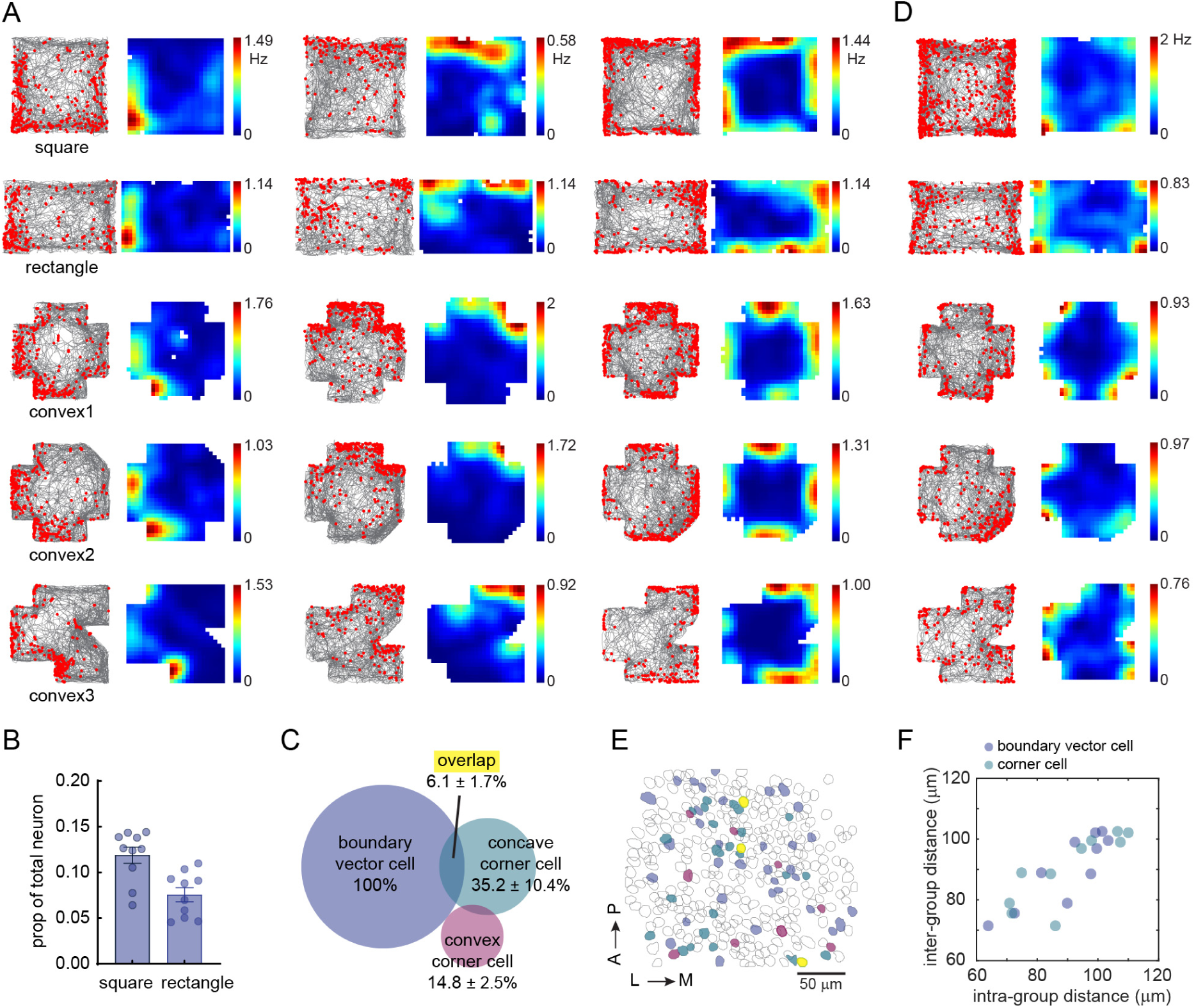
Corner coding differs from boundary vector coding in the subiculum. **(A)** Raster plots and rate maps of three boundary vector cells (BVCs) from three different mice, plotted as in Fig. 1F. BVCs were identified in the square environment. Each column is a cell in which its activity was tracked across sessions. **(B)** Proportion of neurons classified as BVCs in the square and rectangle sessions. Each dot represents a session (n = 10 from 10 mice). Histogram and error bars indicate mean ± SEM. **(C)** Venn diagram showing the overlap between BVCs and concave/convex corner cells. BVCs and concave corner cells were identified in the square environment, while convex corner cells were identified in the convex1 arena. All numbers were normalized to the number of BVCs. **(D)** An example neuron classified as both a BVC and concave corner cell based on its activity in the square environment. **(E)** Anatomical locations of BVCs, concave and convex corner cells from a representative mouse. Color codes are the same as in (C). Unfilled grey circles represent other subicular neurons. A: anterior; P: posterior; L: lateral; M: medial. **(F)** Pairwise intra- vs. inter-group anatomical distances for BVCs and corner cells (concave + convex) (repeated measures ANOVA: p = 0.97; n = 10 mice).

## Discussion

Animals use boundaries and corners to orient themselves during navigation (*1–9*). These features that define the geometry of an environment and can serve as landmarks or indicate locations associated with ethologically relevant needs, such as a nest site or an entryway. Here, we report that in addition to neurons that encode environmental boundaries (*22, 27*), the subiculum contains two distinct populations of neurons that encode concave and convex environmental corners.

One remaining question is how corner specific firing patterns are generated within the context of the larger parahippocampal circuit. Anatomical and physiological works have demonstrated the subiculum contains recurrent connections (*36, 46*). Thus one possibility is that corner cell firing patterns reflect a thresholded sum of input from nearby BVCs (*47*), as both corner and boundary coding are largely invariant across environments (Fig. 1F-J, Fig. S2G-H, Fig. 5A). Alternatively, corner cell firing patterns could reflect convergent inputs from CA1 place cells, given the anatomical observations of dense CA1 to subiculum connectivity (*36*). This idea aligns with previously observed clustering of place fields near the corners of the environment in both individual CA1 place cells and temporally synchronized CA1 ensembles (*40, 48*). It is also possible that both place cells and BVCs contribute to corner coding in the subiculum. Future research using targeted manipulations in the hippocampus could help answer this question (*49*).

Given the subiculum contains relatively discrete populations of neurons for encoding boundaries (BVCs), as well as convex and concave corners, the subiculum may be well positioned to provide information to other brain regions regarding the geometry of the environment in an allocentric reference frame. To guide behavior however, this allocentric information needs to interface with egocentric information regarding an animal’s movements (*33*). One possibility is that corner cells in the subiculum could provide a key input to the recently observed corner-associated activity in the lateral entorhinal cortex (LEC) (*50*). Unlike corner coding in the subiculum, LEC corner-associated activity is egocentric and speed modulated, raising the possibility that LEC integrates allocentric corner information with egocentric and self-motion information to prepare an animal to make appropriate actions when approaching a corner (e.g., deceleration or turning).

## Materials and Methods

### Subjects

All procedures were conducted according to the National Institutes of Health guidelines for animal care and use and approved by the Institutional Animal Care and Use Committee at Stanford University School of Medicine and the University of California, Irvine. For this study, 8 Camk2a-Cre; Ai163 (*39*) mice (4 male and 4 female), 1 Camk2-Cre mouse (female, JAX: 005359), and 1 C57BL/6 mouse (male) were used. For the Camk2-Cre mouse, AAV1-CAG-FLEX-GCaMP7f was injected in the right subiculum at anteroposterior (AP): -3.40 mm; lateromedial (ML): +1.88 mm; and dorsoventral (DV): −1.70 mm. For the C57BL/6 mouse, AAV1-Camk2a-GCaMP6f was injected in the right subiculum at the same coordinates. We combined all mice together for all analyses. Mice were group housed with same-sex littermates until the time of surgery. At the time of surgery, mice were 8 -12 weeks old. After surgery mice were singly housed. Mice were kept on a 12-hour light/dark cycle and had ad libitum access to food and water in their home cages at all times. All experiments were carried out during the light phase.

### GRIN lens implantation and baseplate placement

Mice were anesthetized with continuous 1 – 1.5% isoflurane and head fixed in a rodent stereotax. A three-axis digitally controlled micromanipulator guided by a digital atlas was used to determine bregma and lambda coordinates. To implant the gradient refractive index (GRIN) lens above the subiculum, a 1.8 mm-diameter circular craniotomy was made over the posterior cortex (centered at -3.28 mm anterior/posterior and +2 mm medial/lateral, relative to bregma). The dura was then gently removed and the cortex directly below the craniotomy aspirated using a 27- or 30-gauge blunt syringe needle attached to a vacuum pump under constant irrigation with sterile saline. The aspiration removed the corpus callosum and part of the dorsal hippocampal commissure above the imaging window but left the alveus intact. Excessive bleeding was controlled using a hemostatic sponge that had been torn into small pieces and soaked in sterile saline. The GRIN lens (0.25 pitch, 0.55 NA, 1.8 mm diameter and 4.31 mm in length, Edmund Optics) was then slowly lowered with a stereotaxic arm to the subiculum to a depth of -1.75 mm relative to the measurement of the skull surface at bregma. The GRIN lens was then fixed with cyanoacrylate and dental cement. Kwik-Sil (World Precision Instruments) was used to cover the lens at the end of surgery. Two weeks after the implantation of the GRIN lens, a small aluminum baseplate was cemented to the animal’s head on top of the existing dental cement. Specifically, Kwik-Sil was removed to expose the GRIN lens. A miniscope was then fitted into the baseplate and locked in position so that the GCaMP-expressing neurons and visible landmarks, such as blood vessels, were in focus in the field of view. After the installation of the baseplate, the imaging window was fixed for long-term, in respect to the miniscope used during installation. Thus, each mouse had a dedicated miniscope for all experiments. When not imaging, a plastic cap was placed in the baseplate to protect the GRIN lens from dust and dirt.

### Behavioral experiments with imaging

After mice had fully recovered from the surgery, they were handled and allowed to habituate to wearing the head-mounted miniscope by freely exploring an open arena for 20 minutes every day for one week. The actual experiments took place in a different room from the habituation. The behavior rig, an 80/20 built compartment, in this dedicated room had two white walls and one black wall with salient decorations as distal visual cues, which were kept constant over the course of the entire study. For experiments described below, all the walls of the arenas were acrylic and were tightly wrapped with black paper by default to reduce potential reflections from the LEDs on the scope. A local visual cue was always available on one of the walls in the arena. In each experiment, the floors of the arenas were covered with corn bedding. All animals’ movements were voluntary.

#### Circle, equilateral triangle, square, hexagon and low-wall square

This set of experiments was carried out in a circle, an equilateral triangle, a square, a hexagon, and a low-wall square environment. The diameter of the circle was 35 cm. The side lengths were 30 cm for the equilateral triangle and square, and 18.5 cm for the hexagon. The height of all the environments was 30 cm except for the low-wall square, which was 15 cm. In total, we conducted 15, 18, 17, 18, and 12 sessions (20 min / session) from 9 mice in the circular, triangular, square, hexagonal, and low-wall square arenas, respectively. We recorded a maximum of two sessions per condition per mouse. For each mouse, we recorded 1-2 sessions in each day. If two sessions were made from the same animal in a given day, recordings were carried out from different conditions with at least a two-hour gap between sessions. For each mouse, data from this set of experiments were aligned and concatenated, and the activity of neurons were tracked across the sessions. As described above, all the walls of the arenas were black. A local visual cue (strips of white masking tape) was present on one wall of each arena, covering the top half of the wall.

#### 30-60-90 right triangle and trapezoid

This set of experiments was carried out in a right triangle (30 °, 60 °, 90 °) and a trapezoid environment. Corner angles from the trapezoid were 55°, 90°, 90° and 125°. The dimensions of the mazes were 46 (L) x 28 (W) x 30 (H) cm. In total, we conducted 16 sessions each (25 min / session) from 8 mice for the right triangle and trapezoid. Data from this set of experiments were aligned and concatenated, and the activity of neurons was tracked across the sessions for each mouse. Other recording protocols were the same as described above.

#### Insertion of a discrete corner in a square environment

This set of experiments was carried out in a large square environment with dimensions of 40 (L) x 40 (W) x 40 (H) cm. The experiments comprised a baseline session followed by four sessions with the insertion of a discrete corner into the square maze. In these sessions, the walls that formed the discrete corner were gradually separated by 0, 1.5, 3, and 6 cm. Starting from 3 cm, the animals were able to pass through the gap without difficulty. The dimensions of the inserted walls were 15 (W) x 30 (H) cm. For each condition, we recorded 8 sessions (30 min / session) from 8 mice by conducting a single session from each mouse per day. Data from this set of experiments were aligned and concatenated, and the activity of neurons was tracked throughout the sessions.

#### Square, rectangle, convex-1, convex-2, convex-3, and convex-m1

This set of experiments was carried out in a large square, rectangle, and multiple convex environments that contained both concave and convex corners. The dimensions of the square were 40 (L) x 40 (W) x 40 (H) cm and the rectangle were 46 (L) x 28 (W) x 30 (H) cm. The convex arenas were all constructed based on the square environment using wood blocks or PVC sheets that were tightly wrapped with the same black paper. There convex corners had angles at 270° and 315° in the convex environments. Note that, for 4 out of 10 mice, their convex-2 and -3 arenas were constructed in a mirrored layout compared to the arenas of the other six mice to control for any potential biases that could arise from the specific geometric configurations in the environment (Fig. 4C). For convex-m1 (Fig. S4A), the northeast convex corner was decorated with white, rough surface masking tape from the bottom all the way up to the top of the corner. For each condition, we recorded 10 sessions (30 min / session) from 10 mice, a single session from each mouse per day. For each mouse, data from this set of experiments were aligned and concatenated, and the activity of neurons was tracked across all the sessions.

#### Convex environment with an obtuse convex corner

This set of experiments was carried out in a convex environment that contained two 270° convex corners and one 225° convex corner (Fig. S4D). The arena was constructed in the same manner as the other convex environments described above. For two days, we recorded a total of 18 sessions (30 min / session) from 9 mice, two sessions per mouse. Please note, although the maze was rotated by 90° in the 2^nd^ session, we combined the two sessions together for the analysis.

#### Shuttle box

The shuttle box consisted of two connected, 25 (L) x 25 (W) x 25 (H) cm compartments with distinct colors and visual cues (Fig. S3A). The opening in the middle was 6.5 cm wide, so that the mouse could easily run between the two compartments during miniscope recordings. The black compartment was wrapped in black paper, but not the grey compartment. For two days, we recorded a total of 18 sessions (20 min / session) from 9 mice, two sessions per mouse.

#### Recordings in the dark or with trimmed whiskers

This set of experiments was carried out in a square environment with dimensions of 30 (L) x 30 (W) x 30 (H) cm. The animals had experience in the environment before this experiment. The experiments consisted of three sessions: a baseline session, a session recorded in complete darkness, and a session recorded after the mice’s whiskers were trimmed. For the dark recording, the ambient light was turned off immediately after the animal was placed inside the square box. The red LED (∼650 nm) on the miniscope was covered by black masking tape. This masking did not completely block the red light, so the behavioral camera could still detect the animal’s position. Before the masking, the intensity of the red LED was measured as ∼12 lux from the distance to the animal’s head. However, after the masking, the intensity of the masked LED was comparable to the measurement taken with the light meter sensor blocked (complete darkness, ∼2 lux). The blue LED on the miniscope was completely blocked from outside. For the whisker-trimmed session, facial whiskers were trimmed 12 hours before the recording. For each condition, we recorded 9 sessions (20 min / session) from 9 mice by conducting a single session from each mouse per day. For each mouse, data from this set of experiments were aligned and concatenated, and the activity of neurons was tracked across these sessions.

### Miniscope imaging data acquisition and pre-processing

Technical details for the custom-constructed miniscopes and general processing analyses are described in (*36, 40, 51*) and at miniscope.org. Briefly, this head-mounted scope had a mass of about 3 grams and a single, flexible coaxial cable that carried power, control signals, and imaging data to the miniscope open-source Data Acquisition (DAQ) hardware and software. In our experiments, we used Miniscope V3, which had a 700 μm x 450 μm field of view with a resolution of 752 pixels x 480 pixels (∼1 μm per pixel). Images were acquired at ∼30 frames per second (fps) and recorded to uncompressed avi files. The DAQ software also recorded the simultaneous behavior of the mouse through a high definition webcam (Logitech) at ∼30 fps, with time stamps applied to both video streams for offline alignment.

For each set of experiments, miniscope videos of individual sessions were first concatenated and down-sampled by a factor of 2, then motion corrected using the NoRMCorre MATLAB package (*52*). To align the videos across different sessions for each animal, we applied an automatic 2D image registration method (github.com/fordanic/image-registration) with rigid x-y translations according to the maximum intensity projection images for each session. The registered videos for each animal were then concatenated together in chronological order to generate a combined data set for extracting calcium activity.

To extract the calcium activity from the combined data set, we used extended constrained non-negative matrix factorization for endoscopic data (CNMF-E) (*53, 54*), which enables simultaneous denoising, deconvolving and demixing of calcium imaging data. A key feature includes modeling the large, rapidly fluctuating background, allowing good separation of single-neuron signals from background, and separation of partially overlapping neurons by taking a neuron’s spatial and temporal information into account (see (*53*) for details). A deconvolution algorithm called OASIS (*55*) was then applied to obtain the denoised neural activity and deconvolved spiking activity (Fig. S1B). These extracted calcium signals for the combined data set were then split back into each session according to their individual frame numbers. As the combined data set was large (> 10GB), we used the Sherlock HPC cluster hosted by Stanford University to process the data across 8 – 12 cores and 600 – 700 GB of RAM. While processing this combined data set required significant computing resources, it enhanced our ability to track cells across sessions from different days. This process made it unnecessary to perform individual footprint alignment or cell registration across sessions. The position, head direction and speed of the animals were determined by applying a custom MATLAB script to the animal’s behavioral tracking video. Time points at which the speed of the animal was lower than 2 cm/s were identified and excluded from further analysis. We then used linear interpolation to temporally align the position data to the calcium imaging data.

### Corner cell analyses

#### Calculation of spatial rate maps

After obtained the deconvolved spiking activity of neurons, we binarized it by applying a threshold using a 3 x standard deviation of all the deconvolved spiking activity for each neuron. The position data was sorted into 1.6 x 1.6 cm non-overlapping spatial bins. The spatial rate map for each neuron was constructed by dividing the total number of calcium spikes by the animal’s total occupancy in a given spatial bin. The rate maps were smoothed using a 2D convolution with a Gaussian filter that had a standard deviation of 2.

#### Corner score for each field

To detect spatial fields on a given rate map, we first applied a threshold of 0.3 times the maximum spike rate to the rate map, and identified the location of the field using the x, y coordinates of the maxima for that field. In addition, the coordinates of the centroid and corners of the environments were automatically detected with manual corrections. For each field, we defined the corner score as

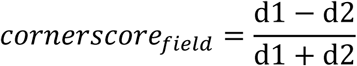

where d1 is the distance between the environmental centroid and the field. d2 is the distance between the field and the nearest environmental corner. The score ranges from -1 for fields situated at the centroid of the arena to +1 for fields perfectly located at a corner (Fig. S1F).

#### Corner score for each cell

There were two situations that needed to be considered when calculating the corner score for each cell (Fig. S1G). First, if a cell had *n* fields in an environment that had *k* corners (*n* ≤ *k*), the corner score for that cell was defined as:

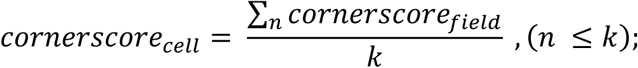

Second, if a cell had more fields than the number of environmental corners (*n* > *k*), the corner score for that cell was defined as the sum of the top *k^th^* corner scores minus the sum of the absolute values of the corner scores for the extra fields, and divided by *k*. Namely,

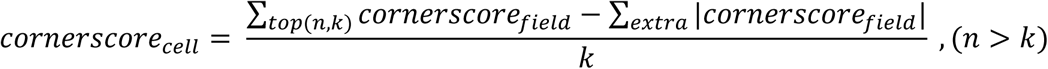

where *top(n,k)* indicates the fields that have the top *k^th^* cornerscore_field_ out of the *n* fields, and ‘extra’ refers to the corner scores for the remaining fields. In this case, the absolute values of the corner scores for the extra fields were used to penalize the final corner score for the cell, so that the score decreased if the cell had too many fields. Note, among all the corner cells identified in the triangle, square and hexagon environments, only 7.4 ± 0.5 % (mean ± SEM; n = 9 mice) of them were classified under this situation, indicating an effective corner score penalization for neurons showing too many fields.

#### Final definition of corner cells

To classify a corner cell, the timing of calcium spikes for each neuron was circularly shuffled 1000 times. For each shuffle, spike times were shifted randomly by 5 - 95% of the total data length, rate maps were regenerated and the corner score for each cell was recalculated. Note, for the recalculation of corner scores for the shuffled rate maps, we did not use the aforementioned penalization process, which resulted in an overall higher shuffled corner score for each neuron. Alternatively, we also attempted to generate the null distribution by shuffling the locations of place fields directly on the original rate map. Although the two methods gave similar results in terms of characterizing corner cells, the latter approach tended to misclassify neurons with few place fields as a corner cell (e.g., a neuron has only one field and the field is in the corner). Therefore, we used the former shuffling method to generate the null distribution. Finally, we defined a corner cell as a cell whose corner score passed the 95^th^ percentile of the shuffled score (Fig. S1H-I), and whose distance between any two fields was greater than half the distance between the corner and centroid of the environment (Fig. S1J).

#### Identification of convex corner cells

To identify convex corner cells, we used similar methods as described above for the concave corner cells, with a minor modification. Namely, after the detection of the field locations on a rate map, we applied a polygon mask to the map using the locations of convex corners as vertices. This polygon mask was generated using the build-in function *poly2mask* in MATLAB. We then considered only the extracted polygon region for calculating corner scores and corresponding shuffles.

#### Measuring the peak spike rate at corners

To measure the peak spike rate at each corner of an environment, we first identified the area near the corner using a 2D convolution between two matrices, M and V. M is the same size as the rate map, containing all zero elements except for the corner bin, which is set to one. V is a square matrix containing elements of ones and can be variable in size. For our analysis, we used a 12 by 12 matrix V, which isolated a corresponding corner region equal to ∼10 cm around the corner. We then took the maximum spike rate in the region as the peak spike rate at the corner. For some specific analyses, due to the unique position or geometry of the corners (e.g., the inserted discrete corner), we decreased the size of the matrix V to obtain a more restricted region of interest for measurement (i.e., ∼5 cm around the corner). To ensure the robustness of our findings, we tried various sizes of the 2D convolution in our analyses, and found that the results were largely consistent with those presented in the manuscript.

#### Corrections of spike rates on the rate map

When comparing spike rates across different corners, it is important to consider the potential impact of the animal’s occupancy and movement patterns on the measurements. To account for any measurements that might have been associated with the animal’s behavior, we generated a simulated rate map using a simulated neuron that fired along the animal’s trajectory using the animal’s measured speed at the overall mean spike rate observed across all neurons of a given mouse. We then used the raw rate map divided by the simulated rate map to obtain the corrected rate map (Fig. S2E). This method ensured that behavior-related factors were present in both the raw and simulated rate maps, and therefore were removed from the corrected rate map.

#### Measuring paired-wise anatomical distances

To measure the pair-wise anatomical distances between neurons, we calculated the Euclidian distance between the centroid locations of each neuron pair under the imaging window for each mouse. We then quantified the average intra-group and inter-group distances for each neuron based on its group identity (e.g., concave vs. convex corner cells). The final result for each group was averaged across all the neurons. We hypothesized that if functionally-defined neuronal groups were anatomically clustered, the inter-group distance would be greater than the intra-group distance.

### Boundary vector cell (BVC) analyses

Rate maps of all the neurons were generated by dividing the open arena into 1.6 cm x 1.6 cm bins and calculating the spike rate in each bin. The maps were smoothed using a 2D convolution with a Gaussian filter that had a standard deviation of 2. To detect boundary vector cells (BVCs), we used a method based on border scores, which we calculated as described previously (*19, 27*):

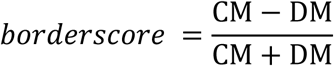

where CM is the proportion of high firing-rate bins located along one of the walls and DM is the normalized mean product of the firing rate and distance of a high firing-rate bin to the nearest wall. We identified BVCs as cells with a border score above 0.6 and whose largest field covered more than 70% of the nearest wall. Additionally, BVCs needed to have significant spatial information (i.e., as in place cells, described below).

### Place cell analyses

#### Spatial information and identification of place cells

To quantify the information content of a given neuron’s activity, we calculated spatial information scores in bits/spike (i.e., calcium spike) for each neuron according to the following formula (*56*),

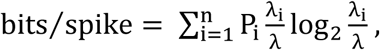

where P_i_ is the probability of the mouse occupying the i-th bin for the neuron, λ_i_ is the neuron’s unsmoothed event rate in the i-th bin, while λ is the mean rate of the neuron across the entire session. Bins with total occupancy time of less than 0.1 second were excluded from the calculation. To identify place cells, the timing of calcium spikes for each neuron was circularly shuffled 1000 times and spatial information (bits/spike) recalculated for each shuffle. This generated a distribution of shuffled information scores for each individual neuron. The value at the 95^th^ % of each shuffled distribution was used as the threshold for classifying a given neuron as a place cell, and we excluded cells with an overall mean spike rate less than the 5^th^ % of the mean spike rate distribution of all the neurons in that animal.

### Position decoding using a naïve Bayes classifier

We used a naïve Bayes classifier to estimate the probability of animal’s location given the activity of all the recorded neurons. The method is described in detail in our previous publication (*40*). Briefly, the binarized, deconvolved spike activity from all neurons was binned into non-overlapping time bins of 0.8 seconds. The M x N spike data matrix, where M is the number of time bins and N is the number of neurons, was then used to train the decoder with an M x 1 vectorized location labels. The posterior probability of observing the animal’s position Y given neural activity X could then be inferred from the Bayes rule as:

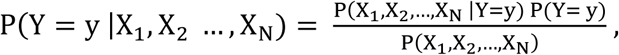

where X = (X_1_, X_2_, … X_N_) is the activity of all neurons, y is one of the spatial bins that the animal visited at a given time, and P(Y = y) is the prior probability of the animal being in spatial bin y. We used an empirical prior as it showed slightly better performance than a flat prior. P(X_1_, X_2_, …, X_N_) is the overall firing probability for all neurons, which can be considered as a constant and does not need to be estimated directly. Thus, the relationship can be simplified to:

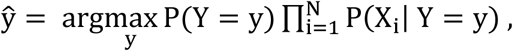

where ŷ is the animal’s predicted location, based on which spatial bin has the maximum probability across all the spatial bins for a given time. To estimate P(X_i_ | Y = y), we applied the built-in function *fitcnb* in MATLAB to fit a multinomial distribution using the bag-of-tokens model with Laplace smoothing.

To reduce occasional erratic jumps in position estimates, we implemented a 2-step Bayesian method by introducing a continuity constraint (*57*), which incorporated information regarding the decoded position in the previous time step and the animal’s running speed to calculate the probability of the current location y. The continuity constraint for all the spatial bins Y at time t followed a 2D gaussian distribution centered at position y_t-1_, which can be written as:

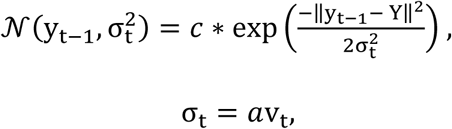

Where *c* is a scaling factor and v_t_ is the instantaneous speed of the animal between time t-1 and t. v_t_ is scaled by *a*, which is empirically selected as 2.5. The final reconstructed position with 2-step Bayesian method can be further written as:

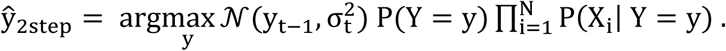

Decoded vectorized positions were then mapped back onto 2D space. The final decoding error was averaged from 10-fold cross-validation. For each fold, the decoding error was calculated as the mean Euclidean distance between the decoded position and the animal’s true position across all time bins.

To test the contribution of corner cells to spatial coding, we first trained the decoder using all neurons and then replaced the neural activity of corner cells with vectors of zeroes from the test data before making predictions. It’s important to note that this activity removal procedure was only applied to the data used for predicting locations and not for training, as ablating neurons directly from the training data will result in the model learning to compensate for the missing information (*58*). We performed this analysis using 10-fold cross-validation for each mouse. To compare the performance of the corner cell removed decoder to the full decoder, we first calculated the 2D decoding error map of a session for each condition, and then obtained a map for error ratio by dividing the error map from the corner cell removed decoder by the error map from the full decoder (Fig. S2C). We then compared the error ratio at the corners of the environment to the center of the environment. For quadrant decoding in the square environment (Fig. S5I-K), we trained and tested the decoder using only the identified corner cells without the 2-step constraint using 10-fold-cross-validation. For the shuffled condition, the decoder was trained and tested for 100 times using circularly shuffled calcium spikes over time. The probability in the correct quadrant was compared between the corner cell trained and shuffled decoders.

### Linear-non-linear Poisson (LN) model

#### Calculation of allocentric and egocentric corner bearing

For each time point in the recording session, the allocentric bearing of the animal to the nearest corner (Fig. S5B) was calculated using the x, y coordinates of the corners and the animal as follows:

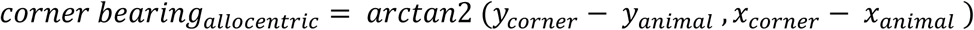

We then derived the egocentric corner bearing of the animal (Fig. S5A-C) by subtracting the animal’s allocentric head direction from the allocentric corner bearing:

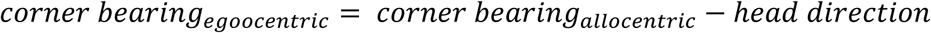

Note that a corner bearing of 0 degrees indicates that the corner was directly in front of the animal, as illustrated in Fig. S5C.

#### Implementation of the linear-non-linear Poisson (LN) model

The LN model is a generalized linear model (GLM) framework which allows unbiased identification of functional cell types encoding multiplexed navigational variables. This framework was described in a previous publication (*59*) and here, we applied the same method to our calcium imaging data in the subiculum. Briefly, 15 models were built in the LN framework, including position (P), head direction (H), speed (S), egocentric corner bearing (E), position & head direction (PH), position & speed (PS), position & egocentric corner bearing (PE), head direction & speed (HS), head direction & egocentric bearing (HE), speed & egocentric bearing (SE), position & head direction & speed (PHS), position & head direction & egocentric bearing (PHE), position & speed & egocentric bearing (PSE), head direction & speed & egocentric bearing (HSE), and position & head direction & speed & egocentric bearing (PHSE). For each model, the dependence of spiking on the corresponding variable(s) was quantified by estimating the spike rate (r_t_) of a neuron during time bin t as an exponential function of the sum of variable values (for example, the animal’s position at time bin t, indicated through an ‘animal-state’ vector) projected onto a corresponding set of parameters (Fig. S5D). This can be mathematically expressed as:

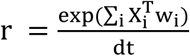

where r is a vector of firing rates for one neuron over T time points, i indexes the variable (i ∈ [P, H, S, E]), X_i_ is the design matrix in which each column is an animal-state vector xi for variable i at one time bin, w_i_ is a column vector of learned parameters that converts animal-state vectors into a firing rate contribution, and dt is the time-bin width.

We used the binarized deconvolved spikes as the neuron spiking data with a time-bin width equal to 500 ms. The design matrix contained the animal’s behavioral state, in which we binned position into 2 cm^2^ bins, head direction and egocentric corner bearing into 20-degree bins, and speed into 2 cm/s bins. Each vector in the design matrix denotes a binned variable value. All elements of this vector are 0, except for a single element that corresponds to the bin of the current animal-state. To learn the variable parameters w_i_, we used the built-in *fminunc* function in MATLAB to maximize the Poisson log-likelihood of the observed spike train (n) given the model spike number (r × dt) and under the prior knowledge that the parameters should be smooth. Model performance for each cell is computed as the increase in Pearson’s correlation (between the predicted and the true firing rates) of the model compared to the 95^th^ % of shuffled correlations (true firing rate was circularly shuffled for 500 times). Performance was quantified through 10-fold cross-validation, where each fold is a random selection of 10% of the data. To determine the best fit model for a given neuron, we used a heuristic forward-search method that determines whether adding variables significantly improved model performance (p < 0.05 for a one-sided sign-rank test, n = 10 cross validation folds).

### Histology

After the imaging experiments were concluded, mice were deeply anesthetized with isoflurane and transcardially perfused with 10 ml of phosphate-buffered saline (PBS), followed by 30 ml of 4% paraformaldehyde-containing phosphate buffer. The brains were removed and left in 4% paraformaldehyde overnight. The next day, samples were transferred to 30% sucrose in PBS and stored in 4°C. At least 24 hours later, the brains were sectioned coronally into 30-µm-thick samples using a microtome (Leica SM2010R, Germany). All sections were counterstained with 10 μM DAPI, mounted and cover-slipped with antifade mounting media (Vectashield). Images were acquired by an automated fluorescent slide scanner (Olympus VS120-S6 slide scanner, Japan) under x10 magnification.

### Data inclusion criteria and statistical analysis

After a certain period post-surgery, the imaging quality began to decline in some animals, and this thus led to slight variations in the number of mice used in each set of experiments, ranging from 8 to 10. We evaluated the imaging quality for each mouse before executing each set of experiments. No mice were excluded from the analyses as long as the experiments were executed. For experiments with two identical sessions for a given condition (e.g., Fig. 1 and 2), sessions with less than 5 identified corner cells were excluded to minimize measurement noise in spike rates. This criterion only resulted in the exclusion of two sessions from two different mice in Fig. 2E.

Analyses and statistical tests were performed using MATLAB (2020a) and GraphPad Prism 9. Data are presented as mean ± SEM. For normality checks, different test methods (D’Agostino & Pearson, Anderson-Darling, Shapiro-Wilk, and Kolmogorov-Smirnov) agreed that almost all data in which we conducted statistical tests followed Gaussian distributions. Thus, a two-tailed paired t-test was used for two-group comparisons throughout the study. For statistical comparisons across more than two groups, repeated measures ANOVA was used prior to pair-wise comparisons. All the statistical tests were performed based on each session or each mouse, as indicated in the corresponding text or figure legend. In all experiments, the level of statistical significance was defined as p ≤ 0.05.

## Acknowledgements

We thank Dr. Zhaoxia Yu for helpful input in designing the method for characterizing corner cells, Sydney Solis for histology assistance, and Dr. Eric Velazquez and Ginny Wu for animal husbandry assistance.

## Author contributions

Conceptualization (YS, DAN, LMG), Methodology (XX, YS), Investigation (YS), Visualization (YS), Funding acquisition (XX, LMG), Project administration (YS, LMG), Supervision (YS, LMG), Writing – original draft (YS, LMG), Writing – review and editing (YS, XX, DAN, LMG).

## Data and materials availability

Upon publication, data will be made available on a publicly accessible data repository site (e.g., figshare, DANDI, Mendeley). Custom scripts for analyzing the data will be available on GitHub.

---

**Fig. S1.**
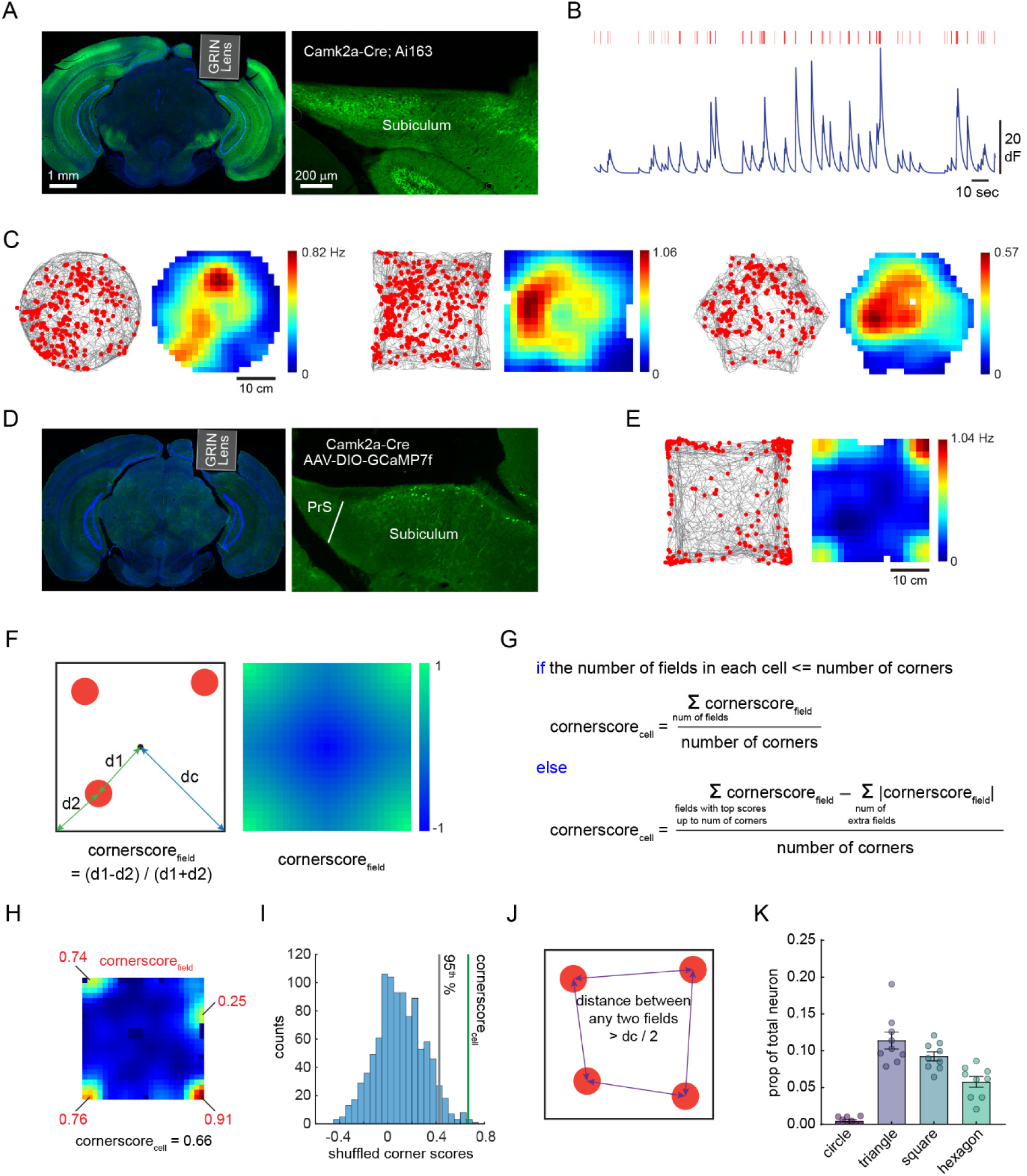
Histology and the development of a score to classify corner cells. **(A)** Histology of GRIN lens implantation in the dorsal subiculum of an example Camk2a-Cre; Ai163 mouse. Green: GCaMP6, Blue: DAPI. Right, enlarged view of GCaMP6-expressing subiculum neurons. **(B)** Representative de-noised calcium signal traces (dark blue) and de-convolved inferred spikes (red bars) extracted from CNMF-E. **(C)** Raster plots and rate maps of a representative place cell in the subiculum. Raster plot (left) indicates extracted spikes (red dots) on top of the animal’s running trajectory (grey lines) and the spatial rate map (right) is color coded for maximum (red) and minimum (blue) values. Activity was tracked across different environments. **(D)** Same as (A), but from a Camk2a-Cre mouse with AAV-DIO-GCaMP7f injected in the subiculum. GCaMP expression was restricted to the subiculum. PrS: presubiculum. **(E)** An example corner cell from the animal in (D), plotted as in (C). **(F)** Left, the definition of corner score for a given spatial field (cornerscore_field_). d1: distance from the center of the arena to the field; d2: distance from the field to the nearest corner; dc: the mean distance from the corners to the center of the arena. Right, the distribution of cornerscore_field_ in a square environment for a given field. Namely, this represents the corner score you would expect if a neuron was active in a given pixel of this plot. Note that the cornerscore can range from -1 (blue) to 1 (green). **(G)** The definition of corner score for a given cell (cornerscore_cell_, see Methods). **(H)** An example corner cell with corner score values for each field labeled in red, and the final corner score for this cell is shown below. **(I)** Shuffling of cornerscore_cell_ to determine a threshold for classifying a neuron as a corner cell. This example is from the same cell in (H). **(J)** An additional constraint for defining a corner cell was that the distance between any two fields needed to be greater than half of the dc value indicated by the blue line in (F). **(K)** same as Fig. 1G, but plotted for each mouse (n = 9 mice) instead of for each session.

**Fig. S2.**
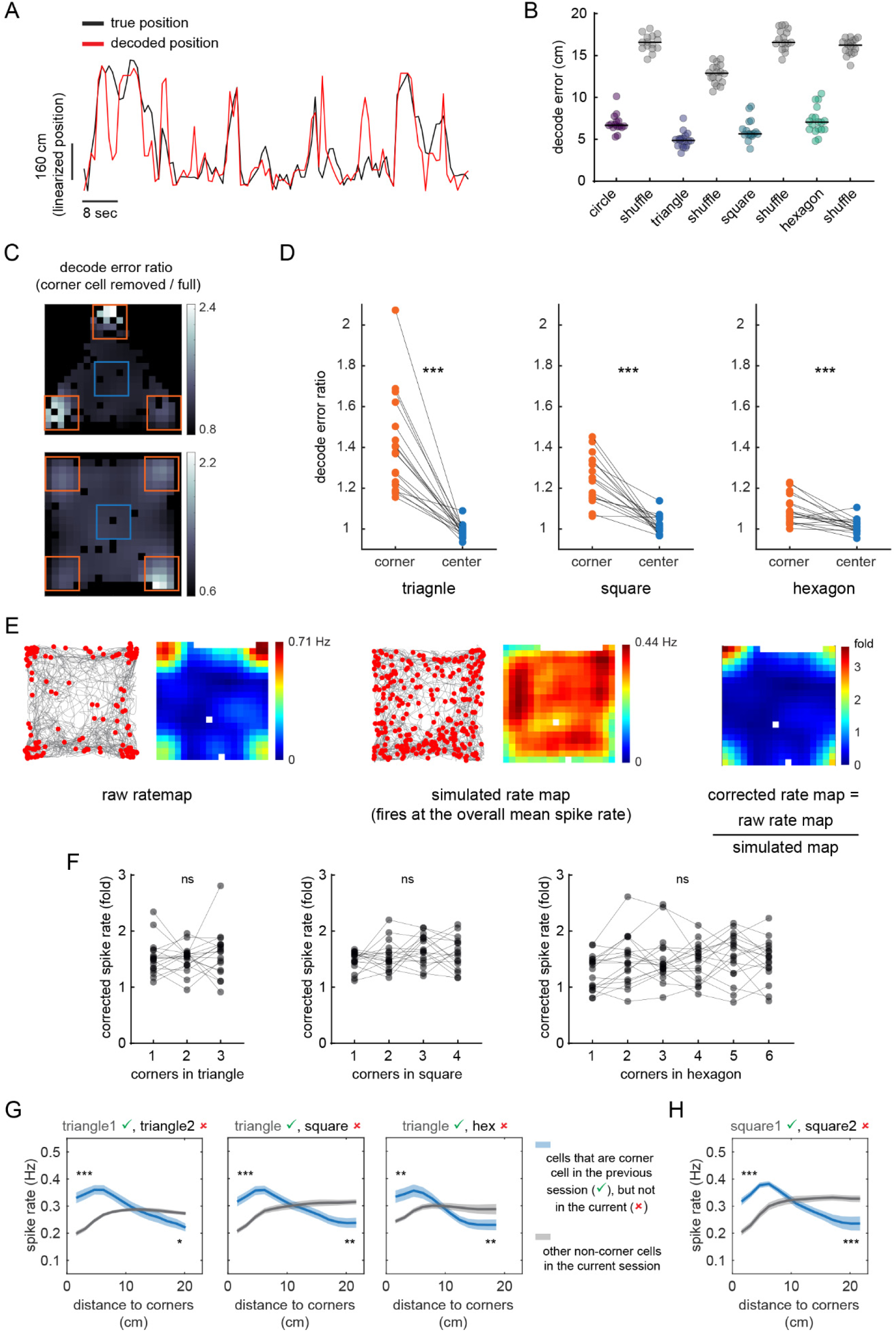
Spatial decoding and peak spike rates of corner cells. **(A)** An example of the true vs. decoded spatial position (linearized) using the full decoder (all the recorded subiculum neurons). **(B)** Decoding performance of the full decoder in different environments. The decoder was trained and tested within each session using 10-fold cross-validation. Each dot is a session, black lines represent the median. Decoding errors were compared with the corresponding shuffle within each condition (two-tailed paired t-test: all p < 0.0001; n = 15, 18, 17, and 18 sessions for circle, triangle, square, and hexagon, respectively, from 9 mice). **(C)** To test the contribution of classified corner cells to spatial coding, we performed another set of decoding analyses in which we removed corner cells from the data before making predictions. Panels indicate the decoding error ratio from a triangle (top) or square (bottom) session, color coded for larger (white) and smaller (black) error ratios. The error ratio was obtained by taking the error map from the corner cell removed decoder and dividing it by the error map of the full decoder. Corner areas (orange boxes) showed higher error ratios than the center area (blue box). **(D)** Quantitative comparisons of the decoding error ratios between the corner and the center areas (shown in C) for all sessions. Each line is a session (two-tailed paired t-test: triangle: t(17) = 7.19, p < 0.0001; square: t(16) = 7.03, p < 0.0001; hexagon: t(17) = 4.47, p = 0.0003; n = 18, 17, and 18 sessions for triangle, square, and hexagon, respectively, from 9 mice). **(E)** Method for obtaining the corrected rate map and calculating the corrected spike rate. For each corrected rate map, a simulated rate map was generated using a simulated neuron that fires along the animal’s trajectory using the animal’s own speed at the overall mean spike rate observed across all neurons of a given mouse (Methods). To obtain the corrected rate map, we divided the raw rate map by the simulated rate map. Therefore, the spike rates on the corrected rate map were automatically converted to fold changes relative to the simulated rate map. This method was used to correct for any measurements that might have been associated with the animal’s movement or occupancy, as purely behavior-related changes should be evident in both the raw and simulated rate maps. **(F)** Corrected peak spike rates of corner cells at the corners of the triangle (n = 18), square (n = 17), and hexagon (n = 18), respectively. Each line represents a session (repeated measures ANOVA: all p > 0.05). ns: not significant. **(G-H)** Spike rates plotted relative to the distance to the nearest corner. Blue curve indicates neurons that are corner cell in the first session (green check mark) but not in the second session (red cross). The plots were generated based on the activity of neurons in the second session. Grey curve indicates other non-corner cells in the second condition. Statistical tests were carried out as in Fig. 1H (two-tailed paired t-test: * p < 0.05, ** p < 0.01, *** p < 0.001; n = 9 sessions from 9 mice for each plot).

**Fig. S3.**
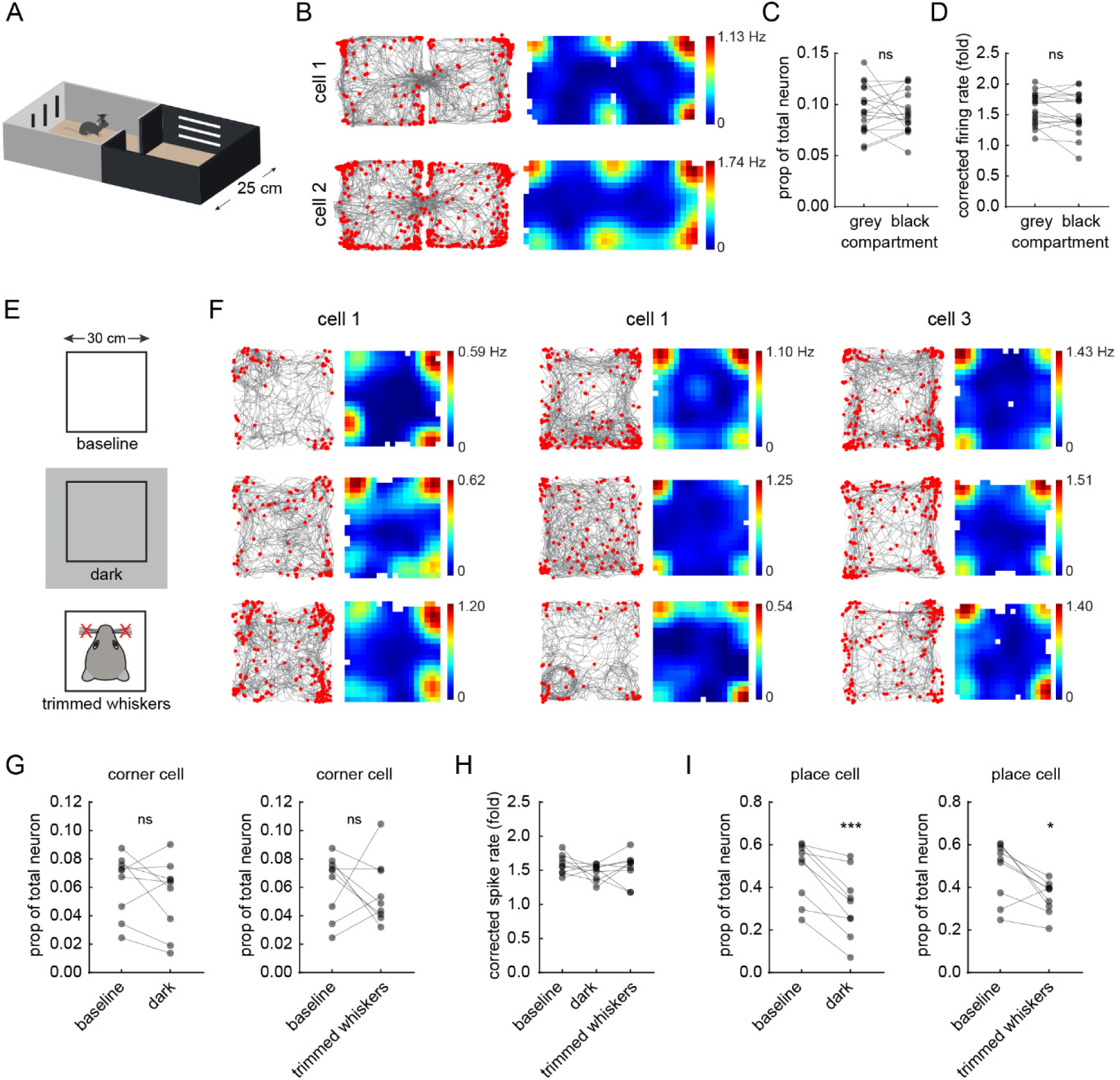
The response of corner cells to non-geometric and sensory manipulations. **(A)** Schematic of a shuttle box composed of two compartments that differed in their visual and tactile cues. **(B)** Two example corner cells from two different mice recorded in the shuttle box shown in (A). Raster plot (left) indicates extracted spikes (red dots) on top of the animal’s running trajectory (grey lines) and the spatial rate map (right) is color coded for maximum (red) and minimum (blue) values. **(C)** Proportion of neurons classified as corner cells in the grey vs. black compartments of the shuttle box (two-tailed paired t-test: t(17) = 0.50, p = 0.62; n = 18 sessions from 9 mice). **(D)** Average corrected peak spike rates of corner cells at the corners in the grey vs. black compartments (two-tailed paired t-test: t(17) = 1.41, p = 0.18, n = 18). **(E)** Schematic of the experiment in which we conducted subiculum recordings in the dark or after trimming of whiskers. **(F)** Raster plots and the corresponding rate maps of three corner cells from three different mice, as in (B). Conditions correspond to the schematic on the left. Each column is a neuron with activity tracked across all the sessions. **(G)** Left, proportion of corner cells between baseline and recordings in the dark (two-tailed paired t-test: t(8) = 1.19, p = 0.27; n = 9 sessions from 9 mice). Right, proportions of corner cells between baseline and recordings after trimming of the whiskers (two-tailed paired t-test: t(8) = 0.53, p = 0.61, n = 9). **(H)** Comparisons of the corrected peak spike rates of corner cells at square corners across all conditions in (E) (repeated measures ANOVA: p = 0.37; n = 9 sessions from 9 mice). **(I)** Same as (G), but for neurons classified as place cells in the subiculum (two-tailed paired t-test: left: t(8) = 5.31, p = 0.0007; right: t(8) = 3.19, p = 0.013; n = 9 sessions from 9 mice). For all panels, ns: not significant.

**Fig. S4.**
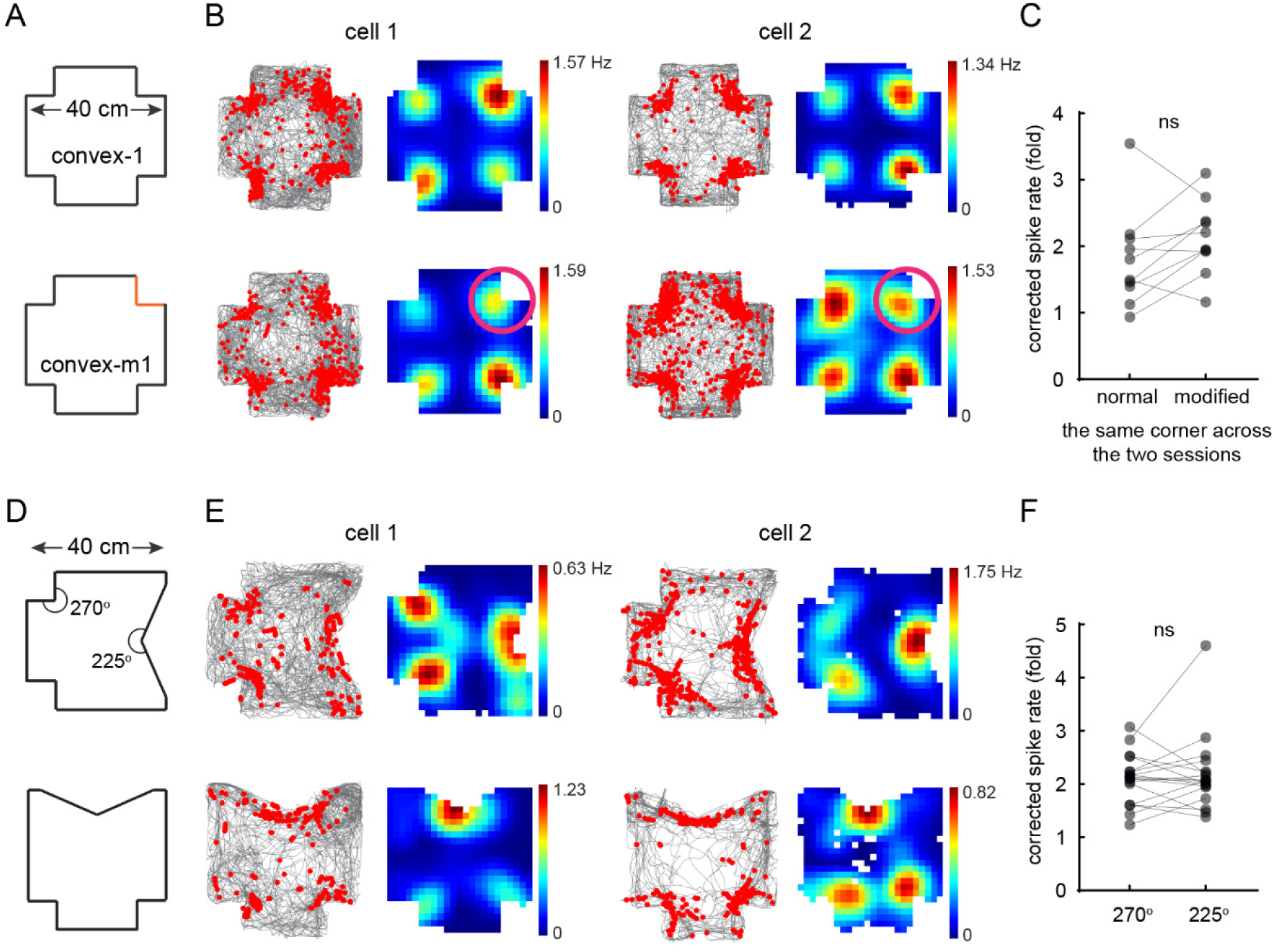
Convex corner cells are not sensitive to non-geometric changes or corner angles. **(A)** Schematic of a normal (convex-1) and modified (convex-m1) convex environments. In convex-m1, one of the convex corners was composed of walls of a different color and texture from the other three. **(B)** Examples of two convex corner cells from two different mice. Raster plot (left) indicates extracted spikes (red dots) on top of the animal’s running trajectory (grey lines) and the spatial rate map (right) is color coded for maximum (red) and minimum (blue) values. Conditions correspond to the schematic on the left. Each column is a neuron in which its activity was tracked across the two conditions. Pink circles delineate the location of the modified corner in the convex-m1 arena. **(C)** Corrected peak spike rates of convex corner cells at the location of the modified corner in the convex-1 vs. convex-m1 arenas (two-tailed paired t-test: t(9) = 1.81, p = 0.10; n = 10 sessions from 10 mice). Corner cells were defined in each session. **(D)** Schematic of a convex environment that contained 270° and 225° corners. The second session was also rotated 90 degrees counterclockwise, but was combined with the first session for analysis. **(E)** Raster plots and the corresponding rate maps of two convex corner cells from two different mice, as in (B). Each column is a neuron in which its activity was tracked across the two sessions. **(F)** Corrected peak spike rates of convex corner cells at 270 ° and 225 ° corners (two-tailed paired t-test: t(17) = 0.67, p = 0.51; n = 18 sessions from 9 mice).

**Fig. S5.**
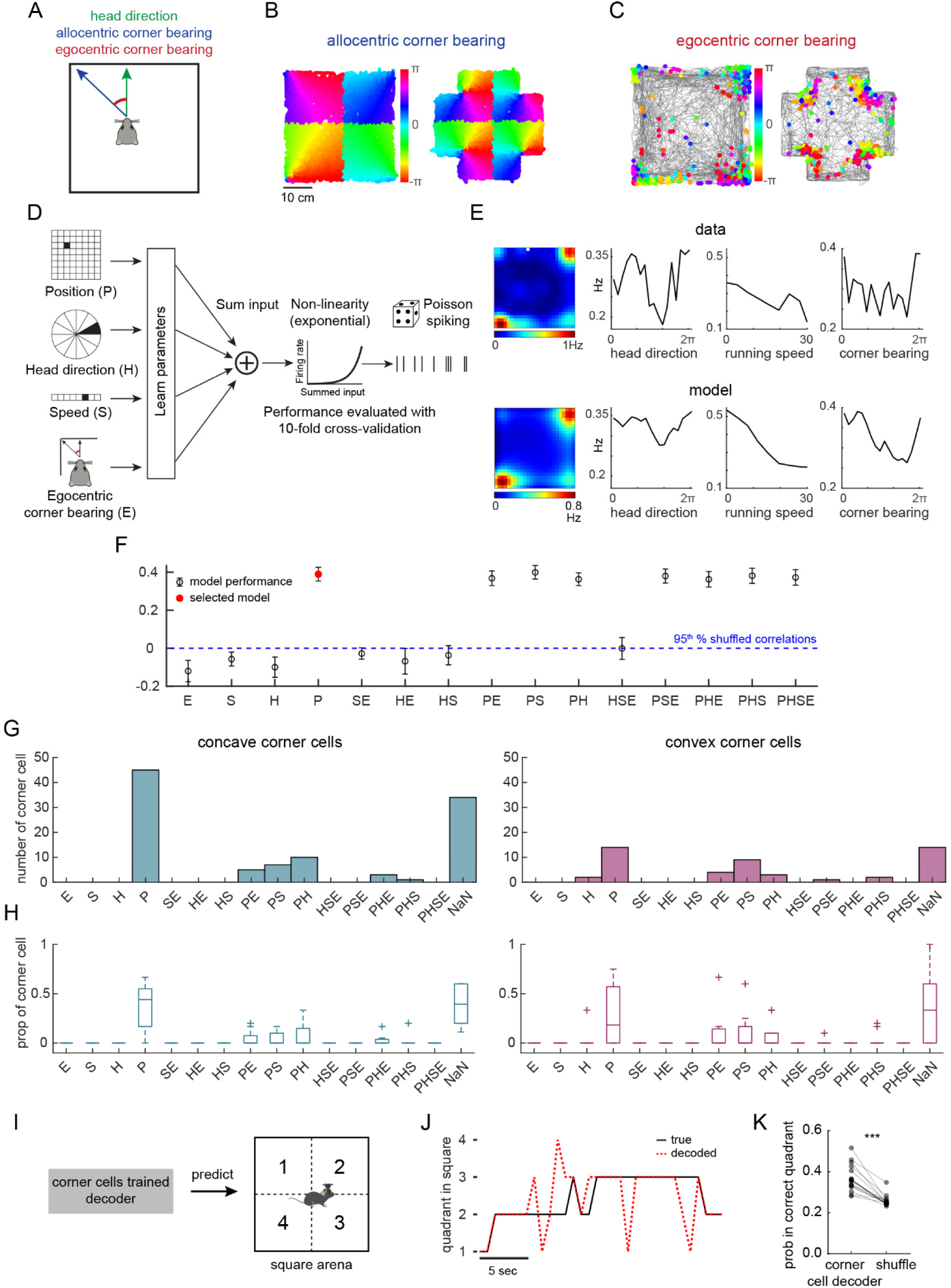
Corner cells primarily correspond to an allocentric reference frame. **(A)** Schematic for calculating egocentric corner bearing (red) by using head direction (green) and allocentric corner bearing (blue) (Methods). **(B)** Examples of allocentric corner bearing in the square and convex-1 environments. **(C)** Corner cell examples with spikes color coded according to the egocentric corner bearing in the square and convex-1 environments. 0 degrees indicates the animal is directly facing to the nearest corner. **(D)** Schematic of the linear-non-linear Poisson (LN) model framework (Methods). **(E)** True tuning curves (top) and model-derived response profiles (bottom) from an example corner cell. **(F)** An example of evaluating the model performance and selecting the best model using a forward search method. This example is from the corner cell in (E) and the best fit model (red dot) is the position (P) only model. **(G)** Number of concave and convex corner cells that fell into each cell type category, respectively. This plot combined all the concave or convex corner cells from all mice identified from the large square or convex 1 in Fig. 4 (a total of 105 concave corner cells and 49 convex corner cells from 10 mice). Some corner cells could not be classified by the model (NaN), potentially due to low spike rates. **(H)** Same as (G), but plotted using the proportion of total corner cells in each animal. For the box plots, the center indicates median, and the box indicates 25th and 75th percentiles. The whiskers extend to the most extreme data points without outliers (+). **(I)** Schematic of using a corner cells trained decoder to predict the animal’s quadrant location in the square arena. **(J)** A decoding trace example of animal’s quadrant location over time using the corner cells trained decoder. Black line: true quadrant location; red dotted line: decoded quadrant location. **(K)** The corner cells trained decoder (mean ± SEM: 0.37 ± 0.02) can predict the animal’s quadrant location more accurately than the shuffle (0.26 ± 0.01) (two-tailed paired t-test: t(16) = 7.97, p < 0.0001; n = 17 sessions from 9 mice).

